# Refining the root-associated microbial consortia for enhanced biocontrol of the root-rot pathogen of corn

**DOI:** 10.1101/2025.09.26.678868

**Authors:** P. Kaki, D. Schlatter, D. Khokhani

## Abstract

Microbial consortia play a crucial role in plant protection by suppressing soil-borne pathogens. A previously studied root-associated microbial consortium consisting of seven bacterial strains (C7) demonstrated biocontrol activity against seedling blight in corn caused by *Fusarium verticillioides*. To enhance its biocontrol potential, we incorporated a free-living bacterial strain (S8) exhibiting biocontrol activity, forming a modified community (C8). We evaluated the biocontrol efficacy of S8, C7, and C8 against four major corn pathogens: *Pythium torulosum, Fusarium graminearum, Fusarium subglutinans*, and *Rhizoctonia solani*. Plate assays revealed that S8 and C8 exhibited the highest inhibition against *P. torulosum* (>65% growth inhibition) but were less effective against *Fusarium* species (25–30%), while none of the communities restricted *R. solani* growth. In pot assays under growth chamber conditions S8 alone exhibited superior pathogen suppression compared to C7 and C8. However, integrating S8 into C7 did not enhance overall biocontrol efficacy. Community analysis via 16S amplicon sequencing revealed no significant shifts in C7 community strain abundance upon S8 introduction, suggesting a lack of establishment of S8 into the C7 community. Given that some individual strains exhibited stronger pathogen suppression than C7 and had variable effects on root biomass and plant height, we designed three sub-communities (SC1, SC2, SC3) based on the highest inhibitory activity. Plant assays demonstrated that inoculation with SC1 and SC2 restored plant height and root biomass, indicating that biocontrol efficacy is primarily driven by specific strain combinations rather than the broader community. Our findings underscore the importance of refining microbial consortia to maximize synergistic interactions and minimize antagonism, advancing sustainable disease management in agriculture.

## 1. Introduction

Corn (*Zea mays* L.), is a global staple crop that faces significant yield losses from soil-borne pathogens like *Fusarium, Pythium*, and *Rhizoctonia* spp. In 2025, corn accounted for 16.7 billion bushels of total production in the United States (USDA; 2025). However, from 2020 to 2024, the estimated economic losses due to corn diseases in these regions reached an average of 963.4 million bushels in 2024 alone, with major pathogens including *Fusarium* spp., *Pythium* spp., and *Rhizoctonia solani* contributing to these losses (Crop Protection Network, 2022; 2024). Generally, developing resistant cultivars is a long-term approach to controlling soilborne diseases (Chang et al., 2025). However, many diseases are still difficult to control due to the limited availability of genetic resistance within germplasm. As a result, fungicides are widely used as an auxiliary control method (Lee et al., 2012). However, excessive reliance on chemical fungicides poses environmental risks, including soil contamination, off-target effects, and can also reduce soil respiration, alter microbial community composition, and disrupt soil enzyme activity (Zubrod et al., 2019; Wang et al., 2025). To address these concerns, biological control offers an eco-friendly alternative by harnessing beneficial microbes to suppress plant pathogens, thereby reducing reliance on chemical inputs.

Early biocontrol strategies focused on applying single microbial strains, employing mechanisms such as antibiotic production or nutrient competition. While effective under controlled conditions, these individual strains often fail to establish or persist within the complex soil environments and are outcompeted by native microbes, resulting in inconsistent field performance (Trabelsi and Mhamdi, 2013; Alori and Babalola, 2018). To overcome these limitations, recent approaches have shifted towards the use of microbial consortia consisting of multiple microbial species with complementary traits. These consortia enhance functional diversity, redundancy, and broaden disease suppression mechanisms, thus offering greater resilience and reliability in agricultural settings (Santos et al., 2019; Minchev et al., 2021). Microbes associated with plants can exhibit several traits, including siderophore production, antibiotic activity, competition for space and nutrients, and the production of hydrolytic enzymes or quorum quenchers that can directly interfere with the pathogen (Legein et al., 2020). Despite their potential, the effectiveness of consortia can be influenced by host genetics, soil properties, and agricultural practices, which can determine the outcome of plant-microbe and microbe-microbe interactions. For example, corn rhizosphere microbiota composition varies with fertilizer use and soil chemistry, underscoring the need for tailored solutions (Peiffer JA, et al., 2013, Aira et al., 2010).

Through host-driven selection, researchers developed a simplified synthetic bacterial community consisting of seven species—*Enterobacter cloacae, Stenotrophomonas maltophilia, Ochrobactrum pituitosum, Herbaspirillum frisingense, Pseudomonas putida, Curtobacterium pusillum*, and *Chryseobacterium indologenes*—representing three of the four dominant bacterial phyla in the corn root microbiome. This seven-member consortium (C7) demonstrated stable root colonization over time, with all strains collectively required to suppress *Fusarium verticillioides*, the causative agent of corn seedling blight (Niu et al., 2017). The enhanced disease suppression observed in microbial consortia is likely attributed to the integration of multiple biocontrol mechanisms. For instance, (Santhanam et al., 2019) reported that a five-strain bacterial consortium exhibited improved rhizoplane colonization and suppression of sudden wilt disease in *Nicotiana attenuata* compared to individual strains. Such improvements in colonization may result from positive microbial interactions that regulate key biological processes, including biofilm formation, growth, and dispersal. Additionally, microbial interactions within consortia can enhance antimicrobial compound production, further strengthening their biocontrol efficacy (Ola at al., 2013). Expanding these communities with additional plant-growth-promoting rhizobacteria (PGPR) that alter root development, exudation patterns, and microbiome dynamics is promising (Vacheron et al., 2013; Ray et al., 2018; Zuluaga et al., 2021; He et al., 2022). For example, inoculation with *Neorhizobium huautlense* T1-17, a PGPR strain that produces IAA, secretes siderophores, and exhibits ACC deaminase activity, increased the proportion of IAA-producing bacteria in the rhizosphere of Chinese cabbage and radish (Wang et al., 2016). These findings highlight the potential of PGPR inoculants to influence biocontrol efficacy through modulation of the microbiome.

In this study, we investigated the integration of a nitrogen-fixing bacterium, *Azotobacter vinelandii* (S8), a strain with potent antifungal activity, into the C7 consortium. While S8 alone exhibited strong inhibitory activity against *Pythium torulosum* and *Fusarium* species, its integration into C7 (forming C8) did not significantly enhance pathogen suppression. Community analysis using 16S sequencing revealed no significant shifts in the relative abundance of strains within the C7 community upon introduction of S8, suggesting that S8 did not functionally integrate within the root-associated community. Given that certain individual strains demonstrated stronger pathogen suppression than the strains combined in the C7 consortium, we hypothesized that biocontrol efficacy is primarily driven by specific strains within the community, rather than by all the strains collectively. To further optimize the biocontrol potential of consortia, we designed three sub-communities (SC1, SC2, SC3) based on the highest inhibitory activity and other phenotypes observed in the plant assays for individual strains. These sub-communities were created to maximize positive interactions and minimize strains that could contribute to antagonistic effects within the consortium. Plant assays revealed that SC1 and SC2 significantly improved plant height and root biomass under pathogen pressure, suggesting that targeted strain selection may be more effective than maintaining a more diverse consortium. Our findings underscore the importance of refining microbial consortia by maximizing positive interactions and minimizing antagonistic effects, ultimately enhancing sustainable disease management in agricultural systems.

## 2. Materials and Methods

### 2.1 Plant, bacterial, and fungal material

Corn (*Zea mays* L.) seeds of the B73 genotype were procured from the Agronomy and Plant Genetics Department of the University of Minnesota Twin Cities (UMN). The seeds were surface sterilized by dipping in 70% (vol/vol) ethanol for 3 min, followed by 5% sodium hypochlorite (vol/vol) for another 3 min, and then rinsed with sterile distilled water. The seven bacterial species (labeled S1 to S7) collected from Roberto Kolter’s laboratory (Niu et al., 2017) were cultured on Tryptic Soy Agar (TSA) medium and incubated at 30□ for 24-48 h. *A. vinelandii* wild-type DJ, called S8, was aerobically grown on Burks medium at 30□. We acquired corn pathogen cultures of *Fusarium graminearum, Fusarium subglutinans, Pythium torulosum*, and *Rhizoctonia solanii* from the Department of Plant Pathology at the University of Minnesota.

### 2.2. *In-vitro* antagonistic activity of bacterial biocontrol agents against corn pathogens

This experiment was performed using Potato Dextrose Agar (PDA) medium, where corn pathogens, including *F. graminearum, F. subglutinans, P. torulosum*, and *R. solanii* were cultured on fresh PDA plates by placing a 1 cm fungal or oomycete plug, obtained using a size 2 cork borer, onto the PDA medium and incubated at 28°C until the mycelium covered the whole plate. Bacterial species (labeled S1 to S7) were streaked on 0.1X TSA medium and incubated at 30°C for 24–48 h. Following incubation, a single colony from each bacterial strain was inoculated into 5 ml of Tryptic Soy Broth (TSB) and incubated overnight at 30°C with shaking at 200 rpm. Subsequently, 50 µl of the overnight culture was transferred to 5 ml of fresh TSB and shaken for an additional 8 h at 30°C. The bacterial cells were harvested by centrifugation and resuspended in 1X PBS buffer. The cell suspensions of each strain were diluted to approximately 10□ cells per milliliter. The mixed communities were obtained from these diluted cultures, where individual cultures were mixed in equal volumes to create a multiple-species bacterial suspension, i.e., C7 and C8 communities. For treatment plates, 20 µl of bacterial suspension was streaked in the center of the plates, and 20 µl of sterile water was used for control treatments. The plates were then incubated at 30°C for 24 h to allow any antimicrobial compounds to diffuse into the medium. After incubation, pathogen plugs of 1 cm were placed perpendicular to the bacterial growth using a size 2 cork borer, and the radial growth of the mycelium was measured seven days post-incubation (dpi).

### 2.3 Growth chamber assays for evaluating the antagonistic activity against the corn pathogens

This experiment was conducted in the growth chambers (Conviron CMP3244; Controlled Environments Ltd., Winnipeg, MB, Canada) located in Alderman Hall at the University of Minnesota. Treatments included S1+Pathogen, S2+Pathogen, S3+Pathogen, S4+Pathogen, S5+Pathogen, S6+Pathogen, S7+Pathogen, S8+Pathogen, and also maintained communities C7+Pathogen, C8+Pathogen, encompassing S1 to S7 and S1 to S8 without pathogen respectively. Furthermore, positive controls consisted of corn plants inoculated solely with the pathogen, while the negative control involved plants without any microbial inoculation. Growth chamber conditions were set to 25 □ during the day for 16 h and 20 °C at night for 8 h with 50% relative humidity. The inocula of the pathogens *F. graminearum, F. subglutinans, P. torulosum*, and *R. solanii* were produced using the sterilized sorghum seeds (Malvick and Bussey, 2010). Briefly, sorghum seeds were soaked overnight in water, drained, and then autoclaved for one hour on two consecutive days. Once the sorghum seeds cooled to room temperature, approximately half of a fully pathogen-colonized PDA plate was added to the autoclaved sorghum seeds using sterile forceps. Then, this mixture was incubated at 28 □ for 14 days with daily shaking to ensure proper aeration. For the plant experiment, sterilized corn seeds were soaked in 50 ml of bacterial inoculum (108 CFU/ml) for respective treatments, and sterile water was used for the control treatment. Each cone was filled to three-quarters with SunGro professional potting mixture (369820SC), followed by a sprinkle of 20 g pathogen-infested sorghum seeds. A 2 cm layer of soil was then added, after which corn seeds were soaked in bacterial inoculum and placed on top of it. Finally, another 2 cm layer of soil was added. These cones were watered daily. Plant growth parameters, including plant height (cm) and fresh root biomass (g), were measured for all treatments at 14 days post-infection with the microbes.

### 2.4 Testing the supernatants of biocontrol agents against *P. torulosum*

Bacterial strains, S1 to S7, were streaked on 0.1X TSA medium and incubated at 30°C for 24–48 h. Following incubation, a single colony from each strain was added to 15 ml of TSB liquid medium and incubated again. The resulting cell suspensions were diluted to an approximate concentration of 10□ CFU/ml for each bacterial strain. For the synthetic communities, 1 mL of each bacterial suspension was combined, followed by incubation in TSB liquid medium for an additional 48 h to allow for any microbial interactions. After this incubation, the mixed suspension was centrifuged, and the supernatant was filtered using a 0.22 µm filter. Subsequently, 20 µl of the resulting cell-free extracts were applied to the center of the treatment plates, while 20 µl of sterile water was used for the control plates. All plates were incubated at 30°C for 24 h to facilitate the diffusion of any antimicrobial compounds into the medium. After incubation, oomycete plugs were placed perpendicular to the streaked supernatant using a size 2 cork borer, and the radial growth of the oomycete mycelium was measured 7 days post-incubation.

### 2.5 DNA extraction and PCR amplification

Bacterial strains were streaked onto appropriate medium plates and incubated at 30°C for 24–48h. A single colony from each strain was inoculated into 5 ml of TSB and incubated overnight at 30°C with constant shaking of 120-125 rpm. Subsequently, 50 µl of the overnight culture was transferred into 5 ml of fresh TSB and incubated under the same conditions for an additional 24– 48 h. Bacterial cells were harvested by centrifugation and resuspended in a 1× PBS buffer. The cell suspensions were adjusted to a final concentration of ∼10□ CFU/mL, and equal volumes of each strain were combined to form the multi-species bacterial communities (C7 and C8). The prepared bacterial communities were incubated at three time points, 5, 10, and 15 days, and genomic DNA was extracted using the FastDNA SPIN Kit for Soil (MP Biomedicals) following the manufacturer’s instructions. All DNA samples were subsequently diluted to a final concentration of 20 ng/µl and used as templates for PCR. Each sample was amplified using the primers F515 (5′-GTGCCAGCMGCCGCGGTAA-3′) and R806 (5′-GGACTACHVGGGTWTCTAAT-3′) targeting the V4 region of the 16S rRNA gene. The extracted DNA was submitted to the University of Minnesota Genomic Center (UMGC) for amplification and sequencing using established protocols (Gohl et al., 2016)

### 2.6 Sequence analysis

Raw sequencing reads were processed using the DADA2 pipeline (v1.26.0; Callahan et al., 2016) in R to generate an amplicon sequence variant (ASV) table. Paired-end reads were quality-filtered, with truncation at 280 bp (forward) and 220 bp (reverse), removal of ambiguous bases, and denoising based on learned error rates. Merged sequences were used to construct an ASV table, followed by chimera removal. 16S sequences were extracted from the S1-S8 genomes using EZbiocloud (Chalita et al., 2024) and used to construct a BLAST database. ASV sequences were then blasted to this database to identify ASVs with 100% identity to these species (>97% of all sequences). Non-target sequences were filtered from the ASV table prior to analyses. Permutational multivariate analysis of variance (PERMANOVA) was used to compare community structure between C7 and C8 at each time point. All analyses were conducted in R (v4.3.1) using the phyloseq, ggplot2, and vegan packages (Niu et al., 2017; McMurdie & Holmes, 2013).

### 2.7 Constructing subcommunities

We systematically compared the inhibitory activity of all individual bacterial strains (S1 to S7) and community (C7) using plate assay, growth chamber conditions, and supernatant tests (Fig. S2). The comparison revealed a varying level of biocontrol activity among the single-species biocontrol agents and the community (C7). Interestingly, we found that individual strains performed better than C7. The previous study (Niu et al., 2017) shows that removal of the S4 strain from the community in a leave-one-out experiment quickly reduced populations of other bacterial strains in the community, suggesting that there might be positive and negative interactions among the community members. Considering the highest inhibitory activity of biocontrol agents, which resulted from the plate, plant and supernatant assay, we addressed the possibility of positive interactions by creating subcommunities. These sub-communities are composed as follows: SC1 (S4, S6, S7), SC2 (S4, S1, S5), and SC3 (S1, S4, S5, S6, S7).

### 2.8 Biochemical assays: IAA and siderophore test

IAA production was measured using the Salkowski reagent assay. The reagent was prepared by dissolving 2.03 g FeCl□ in 25 mL of deionized water (0.5 M stock solution) and mixing 2 mL of this solution with 98 mL of 35% perchloric acid (HClO□). The prepared Salkowski reagent was stored in a sealed container at room temperature until its use. For the assay, 1 mL of culture supernatant was transferred into a sterile 5 mL tube, followed by the addition of 2 mL Salkowski reagent. Samples were incubated at room temperature for 30 min in the dark to allow color development. A blank control was prepared by mixing 1 mL liquid medium with 2 mL Salkowski reagent. Absorbance of the test and blank samples was measured at 530 nm using a spectrophotometer. IAA concentrations (µg/mL) were determined using a standard curve generated from pure IAA (Gang et al., 2019).

Siderophore production was evaluated using the Chrome Azurol S (CAS) agar assay (Louden et al., 2011). All glassware was pre-cleaned with 0.5 M HCl, rinsed with Milli-Q water, and dried before use. The CAS blue dye was prepared by dissolving 0.06 g CAS in 50 mL deionized water (solution 1), 0.0027 g FeCl□·6H□O in 10 mL of 10 mM HCl (solution 2), and 0.073 g HDTMA in 40 mL deionized water (solution 3), which were then combined, autoclaved (liquid cycle, 30 min), and stored until use. Separately, LB agar (pH adjusted to 6.7) was prepared with 100 mL less water than the standard recipe, autoclaved, and maintained in a 60 °C water bath. To prepare CAS agar, 100 mL of the sterile blue dye solution was mixed with molten LB agar, swirled, and poured into Petri plates, which transitioned from yellow to blue upon cooling. For inoculation, bacterial strains (S1–S8) and synthetic communities (SC1, SC2, SC3, C7, and C8) were pre-cultured in tryptic soy broth for 2 days, adjusted to ∼10□ CFU/mL, and 10 µL of each culture was added to the well onto CAS agar plates. The well was created using the downward end of a sterile 1 mL pipette tip. After air-drying in a biosafety cabinet (∼20 min), plates were incubated at 26 °C for 7 days. Siderophore production was indicated by the development of orange or yellow halos against the blue background of the agar.

## 3 Results

### 3.1 Biocontrol activity of the single bacterial strain *Azotobacter vinelandii* (S8) against corn pathogens

We tested the inhibitory activity of the free-living nitrogen-fixing bacterium *Azotobacter vinelandii (S8)* against prominent corn pathogens by performing plate assays and plant experiments in controlled environmental conditions. In plate assays, the inhibitory potential of S8 was evaluated individually against four major corn pathogens: *Rhizoctonia solani* (root rot), *Pythium torulosum* (seedling blight), *Fusarium graminearum* (stalk rot/ear rot), and *Fusarium subglutinans* (stalk rot/ear rot). Among these, S8 exhibited the strongest suppression against *P. torulosum*, followed by moderate inhibition of *F. graminearum* and *F. subglutinans*, while no significant inhibition was observed against *R. solani* (Fig. 1A, B). Based on these results, we performed subsequent growth chamber experiments with *P. torulosum*. In this assay, corn seeds were pre-treated with S8 and sown in soils inoculated with *P. torulosum*. At fourteen days post-inoculation (dpi), S8 treatment significantly reduced the root rot symptoms. We observed improved plant height and root biomass compared to plants infected only with the pathogen or the control plants without any microbial infection (**Fig. 1C, D**). These findings highlight the dual potential of S8 as an effective biocontrol agent against *P. torulosum* and as a plant growth-promoting bacterium.

**Figure 1.**
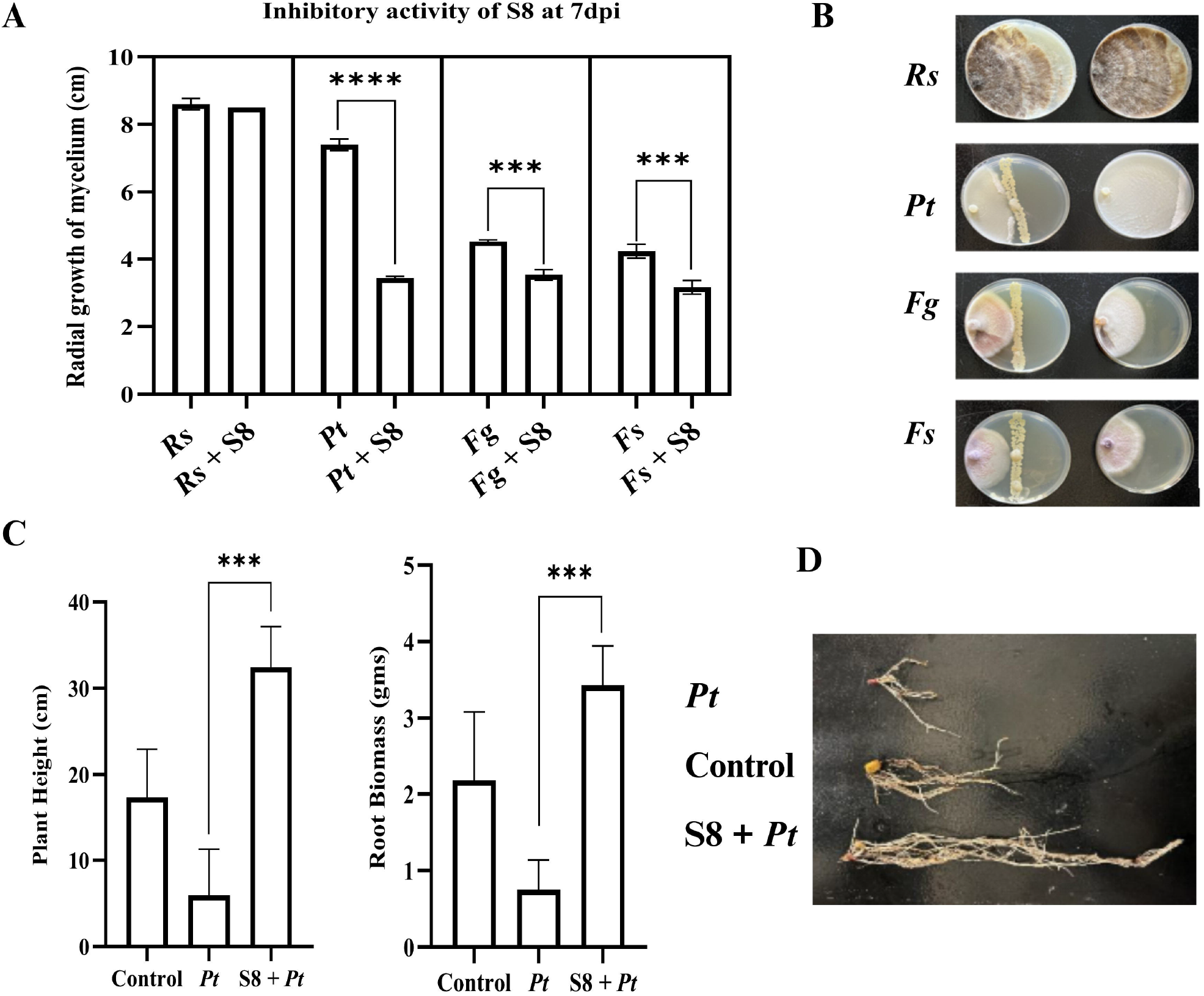
Biocontrol activity of single biocontrol Agent *Azotobacter vinelandii* (S8) against corn fungal and oomycete pathogen. A and B) Plate assay: Measurement of radial growth of corn pathogens namely *Pythium torulosum* (*Pt*), *Fusarium subglutinans* (*Fs*), Fusarium *graminearum* (*Fg*), and *Rhizoctonia solani-AG IIIB* (*Rs*) in cm at 7 days post incubation (dpi). C) Pot assay of S8 against *Pythium torulosum* (*Pt*) where plant height and root biomass was measured at 14 days after infecting with biocontrol agents and pathogen. D) Roots of plants infected only with pathogen, plants with pathogen and S8, plants with no microbial infection (control). Three biological replications were maintained with each including technical replication (n = 5) for plate assay and for pot assay (n = 7). Data are presented as mean ± Standard Deviation. For statistical analysis unpaired *t*-test with Welch’s correction.0’1+ 10.0.2) was used. ***indicates p-values <0.001, **** <0.0001

### 3.2 Antagonistic activity of the synthetic community (C7) and individual bacterial strains

We examined the inhibitory effect of a synthetic community (C7) composed of seven bacterial species (*Stenotrophomonas maltophilia, Ochrobactrum pituitosum, Curtobacterium pusillum, Enterobacter cloacae, Pseudomonas putida, Herbaspirillum frisingense*, and *Chryseobacterium indologenes*) representing three of the four most dominant phyla present in corn roots (Niu et al., 2017). Each of these seven species effectively restricted the growth of *Fusarium verticillioides*, the causal agent of corn seedling blight (Niu et al., 2017). We evaluated the antagonistic activity of the C7 community using plate assay conditions against major corn root rot pathogens, including *R. solani, P. torulosum, F. graminearum*, and *F. subglutinans*. The C7 community exhibited strong inhibitory effects against the oomycete pathogen, *P. torulosum*, and exerted moderate but significant inhibition of *F. graminearum* and *F. subglutinans* (**Fig. 2A**).

**Figure 2.**
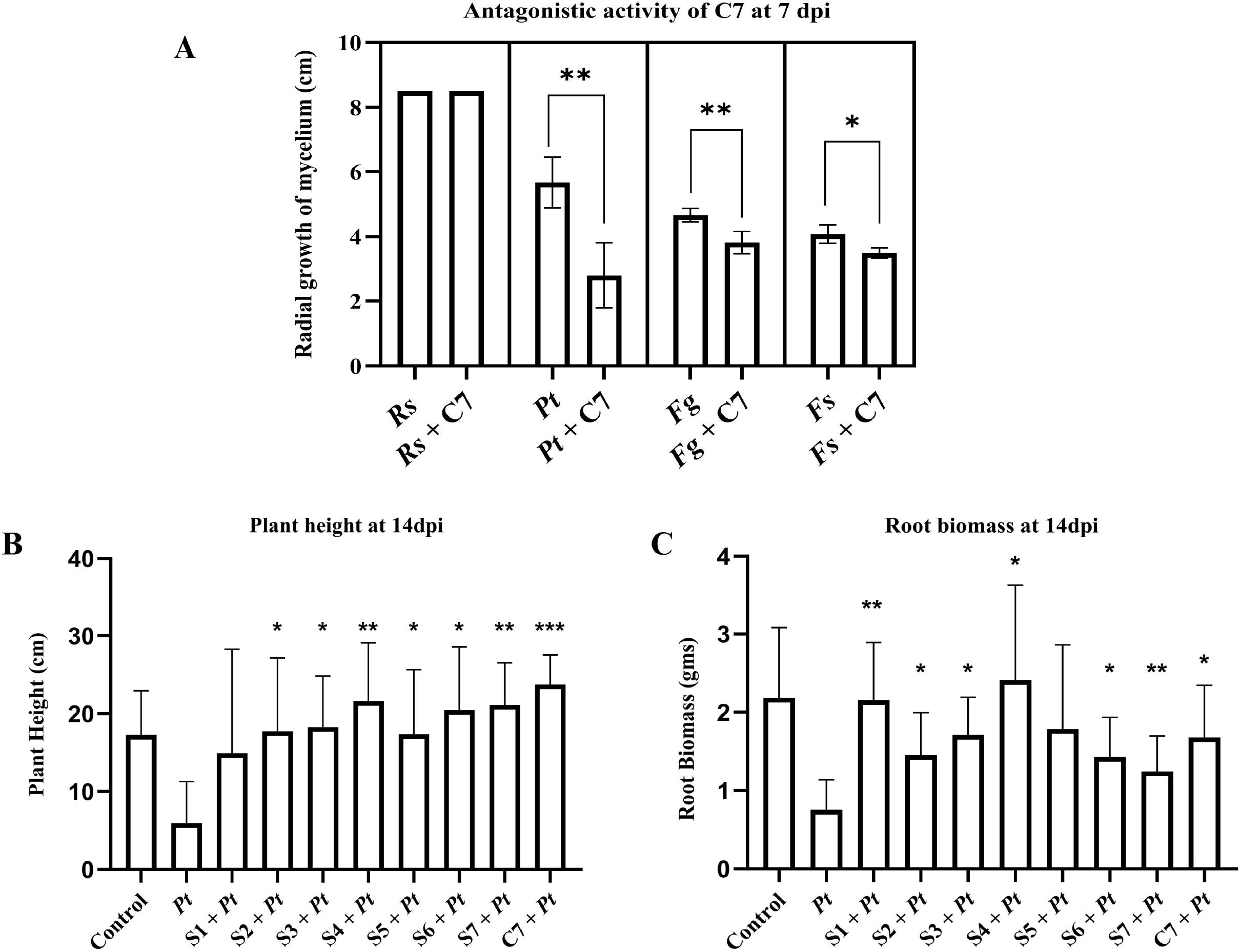
Antagonistic activity of individual bacterial strains (S1 to S7) and community (C7) against corn fungal and oomycete pathogen. A) Plate assay: Measurement of radial growth of corn pathogens in cm at 7 day post incubation (dpi) against *Pythium torulosum* (*Pt*), *Fusarium subglutinans* (*Fs*), *Fusarium graminearum* (*Fg*), and *Rhizoctonia solani* – *AG IIIB* (*Rs*). B-C) Pot assay: Plants were treated with biocontrol strains S1 to S7 or community C7 and inoculated with *Pythium torulosum* (*Pt*). Plant height (B) and root biomass (C) were measured at 14 dpi and compared with plants infected with *pt* alone. Three biological replications were maintained with each including technical replication (n = 5) for plate assay and for pot assay (n = 7). Data are presented as mean ± Standard Deviation. For statistical analysis, unpaired *t*-test with Welch’s correction.0’1+ 10.0.2) was used. * indicates p-values < 0.05, ** < 0.01, *** < 0.001.

To test whether the inhibition activity of C7 translates into protection of corn against the pathogen, we performed plant experiments under controlled growth chamber conditions. We selected *P. torulosum* as a target pathogen since the highest inhibitory effect was seen against it. Plants treated with the C7 community in the presence of *P. torulosum* exhibited greater shoot height compared to treatments with individual bacterial strains and the control treatments. However, certain single strains outperformed the C7 community in promoting root biomass under pathogen pressure (**Fig. 2B and 2C**)

### 3.3 Antagonistic activity of modified community (C8) against corn pathogens under plate and growth chamber conditions

Building on initial screening results, strain S8, identified as the most effective individual antagonist, was incorporated into the existing synthetic community C7 to generate an enhanced microbial consortium, designated as C8. The biocontrol potential of C8 was evaluated against the same set of corn pathogens. *In vitro* assays revealed that C8 exhibited strong inhibition of *P. torulosum* mycelial growth and moderate, yet statistically significant, inhibition of *F. graminearum* and *F. subglutinans* compared to the pathogen-only control. No inhibitory effect was observed against *R. solani*, as treated and control plates displayed comparable growth (**Fig. 3A**). To compare the biocontrol efficacy of individual bacterial strains (S1–S8) and synthetic communities (C7 and C8), we conducted a plate assay against the same set of four corn pathogens. Among all treatments, strain S8 and community C8 exhibited the most potent inhibitory effects, with percent inhibition zones exceeding 65% against *P. torulosum* (**Fig. S1**). Moderate inhibition was observed against *F. subglutinans* and *F. graminearum*, ranging from 25% to 30%, while no detectable inhibition was observed against *R. solani* by any of the individual strains or synthetic communities tested. The C7 community showed a lower inhibition when compared to S8 and C8 against all four corn pathogens. Given this pronounced inhibitory response, we selected *P. torulosum* as the target pathogen for plant experiments. At 14 dpi with the respective biocontrol agents and *P. torulosum*, plants treated with strain S8 alone exhibited the most pronounced recovery in both plant height and root biomass under pathogen pressure, outperforming all other individual strains and both community treatments (**Fig. 3B** and **3C**). In contrast, no significant differences in plant growth parameters were observed between the C7 and C8 communities. The incorporation of S8 into the C7 community did not enhance plant growth promotion or biocontrol efficacy under the tested conditions. This outcome indicates that interactions between S8 and the existing members of C7 may influence the overall performance of the C8 community.

**Figure 3.**
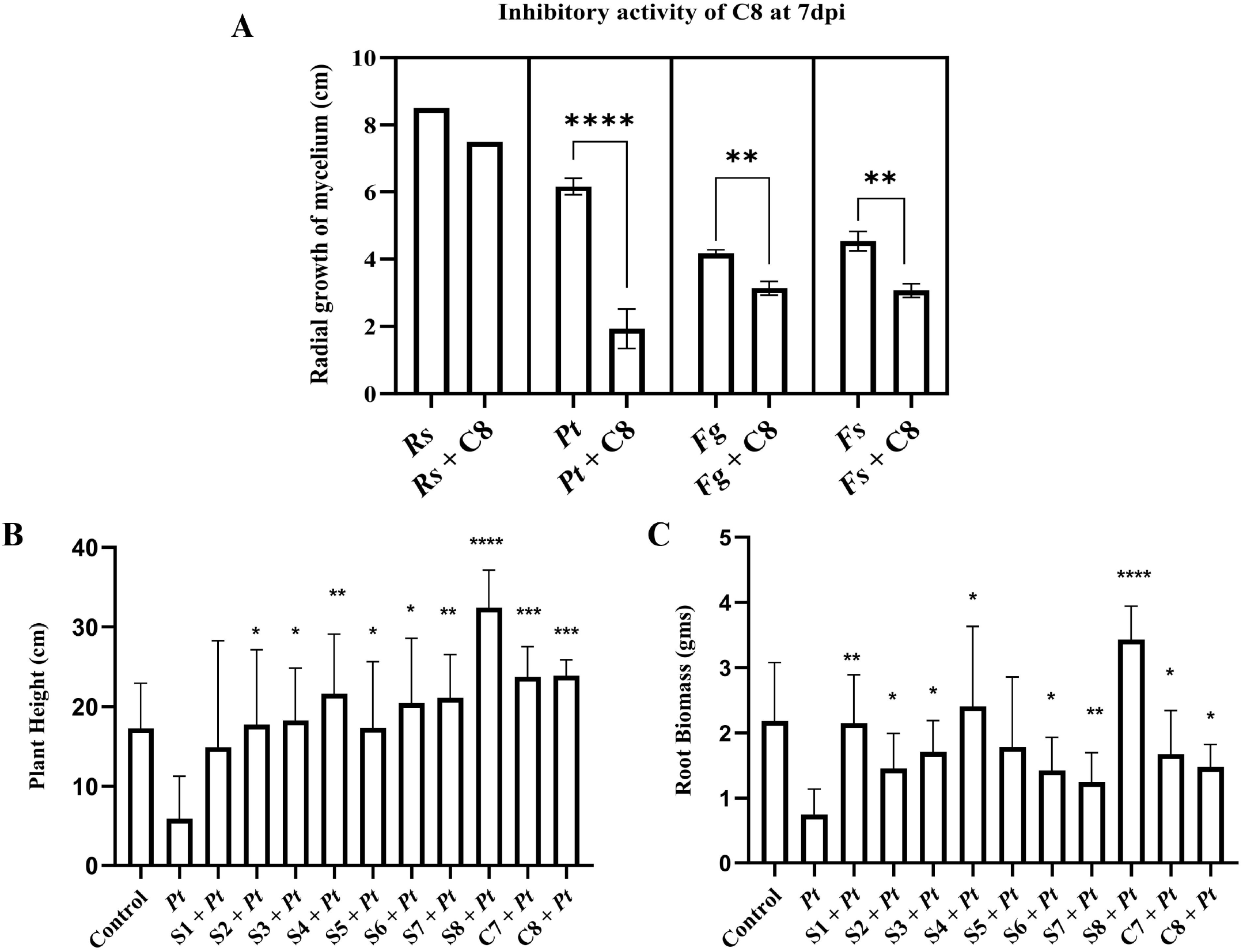
Comparing the biocontrol activity of individual bacterial strains (S1 to S8) and communities (C7 and C8) against the corn pathogen. A) Plate assay: Measurement of radial growth of corn pathogens namely *Pythium torulosum* (*Pt*), *Fusarium subglutinans* (*Fs*), *Fusarium graminearum* (*Fg*), and *Rhizoctonia solani* – *AG IIIB* (*Rs*) in cm at 7 day post incubation (dpi). B-C) Pot assay: Plants were treated with biocontrol strains S1 to S7 or communities C7, C8 and inoculated with *Pythium torulosum* (*Pt*). Plant height (B) and root biomass (C) were measured at 14 dpi and compared with plants infected with *pt* alone. Three biological replications were maintained with each including technical replication (n = 5) for plate assay, for pot assay (n = 7). Data are presented as mean ± Standard Deviation. For statistical analysis unpaired t-test with Welch’s correction.0’1+ 10.0.2) was used. *indicates p values <0.05, ** <0.01, *** <0.001

### 3.4 Temporal dynamics of individual bacterial species in C7 and C8 communities

To assess the population dynamics of the C7 and C8 communities over time, we conducted 16S sequencing at three time points (days 5, 10, and 15) following inoculation. The results showed that the newly added bacterium S8 was detected at a negligible proportion (<0.002%) in the C8 community at any of the time points. Furthermore, no significant differences in communities were observed between the C7 and C8 communities at any time point (**Fig. 4**; PERMANOVA p>0.2 at each time). These findings suggest that the established resilience and robustness of the C7 community may prevent the successful integration of the externally introduced S8 strain, which could explain the lack of enhanced biocontrol efficacy of the C8 community.

**Figure 4.**
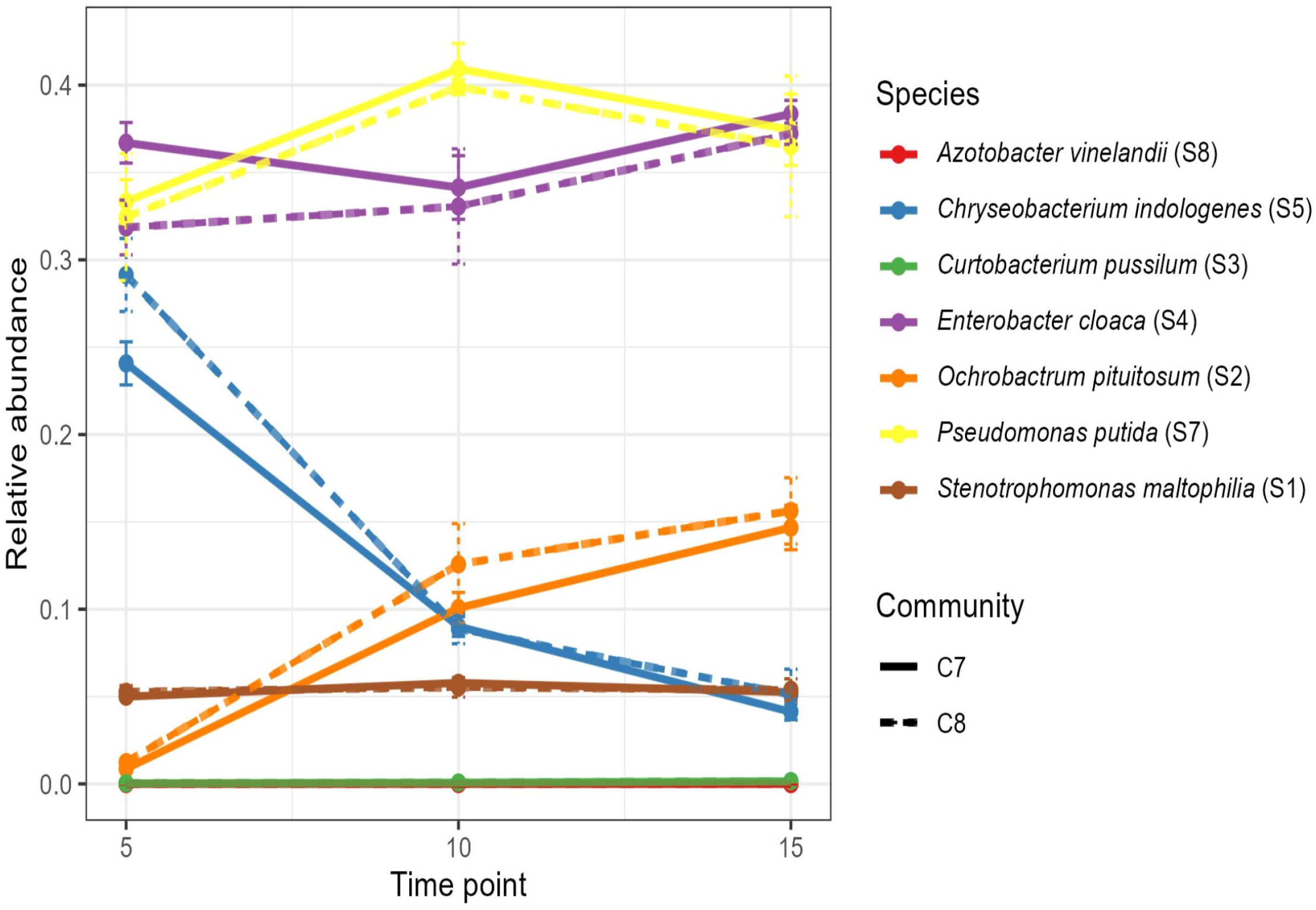
Dynamics of the C7 and C8 bacterial communities were measured using 16S rRNA gene sequencing. Tracked the relative abundance of C7 and C8 communities at three time points (5th, 10th and 15th day). One biological replication was maintained with technical replication (n=3)

### 3.5 Biocontrol and plant growth–promoting traits of individual strains and synthetic communities

#### 3.5.1 Testing Indole-3-acetic acid production

We evaluated indole-3-acetic acid (IAA) production by individual bacterial strains (S1–S8) and synthetic communities (C7 and C8). IAA concentrations differed considerably among strains and across time points. At day 2, strains S2 and S4 produced the highest IAA levels, with S4 reaching ∼75 µg/mL. By day 7, IAA accumulation declined in S2 and S4 but increased in strains such as S3, S5, and S6, which showed higher levels relative to day 2. In contrast, some isolates (e.g., S7 and S8) maintained consistently low IAA levels. Both synthetic communities (C7 and C8) produced moderate and comparable amounts of IAA, with little temporal variation, suggesting that community-level output was more stable compared to the fluctuating responses of individual strains (**Fig. 5A**).

**Figure 5.**
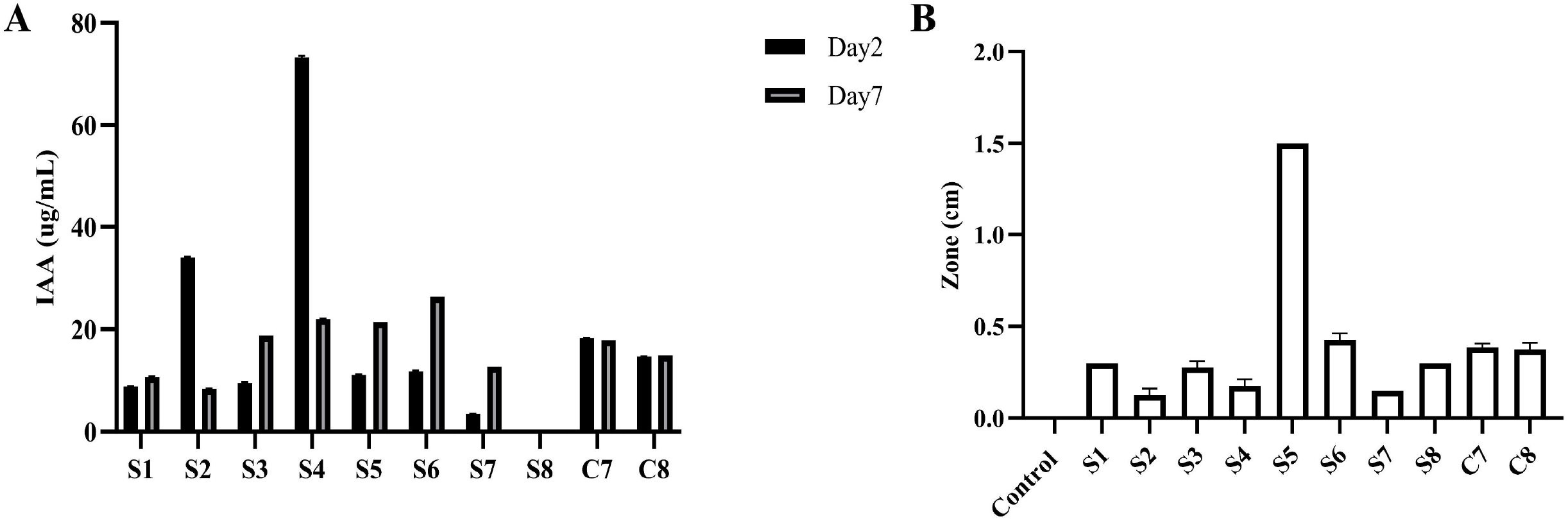
Assessment of IAA and siderophore production by individual strains (S1-S8) and communities (C7 and C8). (A) IAA production was quantified using the Salkowski reagent at 2 and 7 days of incubation. Siderophore production was evaluated by measuring the diameter of the clear zone 7 days post-inoculation. Three biological replications were maintained with technical replications of (n =3). Data are presented as mean ± Standard Deviation.

#### 3.5.2 Testing Siderophore Activity

Siderophore activity, measured as the diameter of the orange halo on CAS agar, also varied among isolates. Strain S5 exhibited the strongest siderophore activity, forming a clear zone of ∼1.5 cm, whereas strains *S1* and S7 showed minimal production. Moderate activity was observed in S6, S8, C7, and C8, with zone diameters around 0.4–0.6 cm, while S2, S3, and S4 displayed weak activity. Control plates lacked detectable siderophore production (**Fig. 5B**). Together, these assays demonstrate that IAA and siderophore production are highly strain-dependent traits. Notably, while strains such as S4 excelled in IAA production and S5 in siderophore activity, no single strain consistently outperformed others across both the traits. The synthetic communities maintained moderate and consistent levels of both metabolites, suggesting a stabilizing effect of community interactions.

### 3.6 Testing the antagonistic activity of sub-communities against the oomycete pathogen under plate and growth chamber conditions

To assess the antagonistic potential of the subcommunities (**Table S1**), we first evaluated their *in vitro* inhibitory activity against *P. torulosum*. All subcommunities displayed inhibition comparable to that of the established biocontrol community C7 (**Fig. 6A**). Subsequently, plant experiments were conducted to evaluate their effects on disease control. Among the subcommunities tested, SC1 (comprising strains S4, S6, and S7) and SC2 (comprising strains S4, S1, and S5) consistently showed enhanced protection from the pathogen and promoted plant growth traits significantly more than both the non-inoculated control and the C7 community. In contrast, SC3 did not confer any measurable benefit (**Fig. 6B, C**, and **D**).

**Figure 6.**
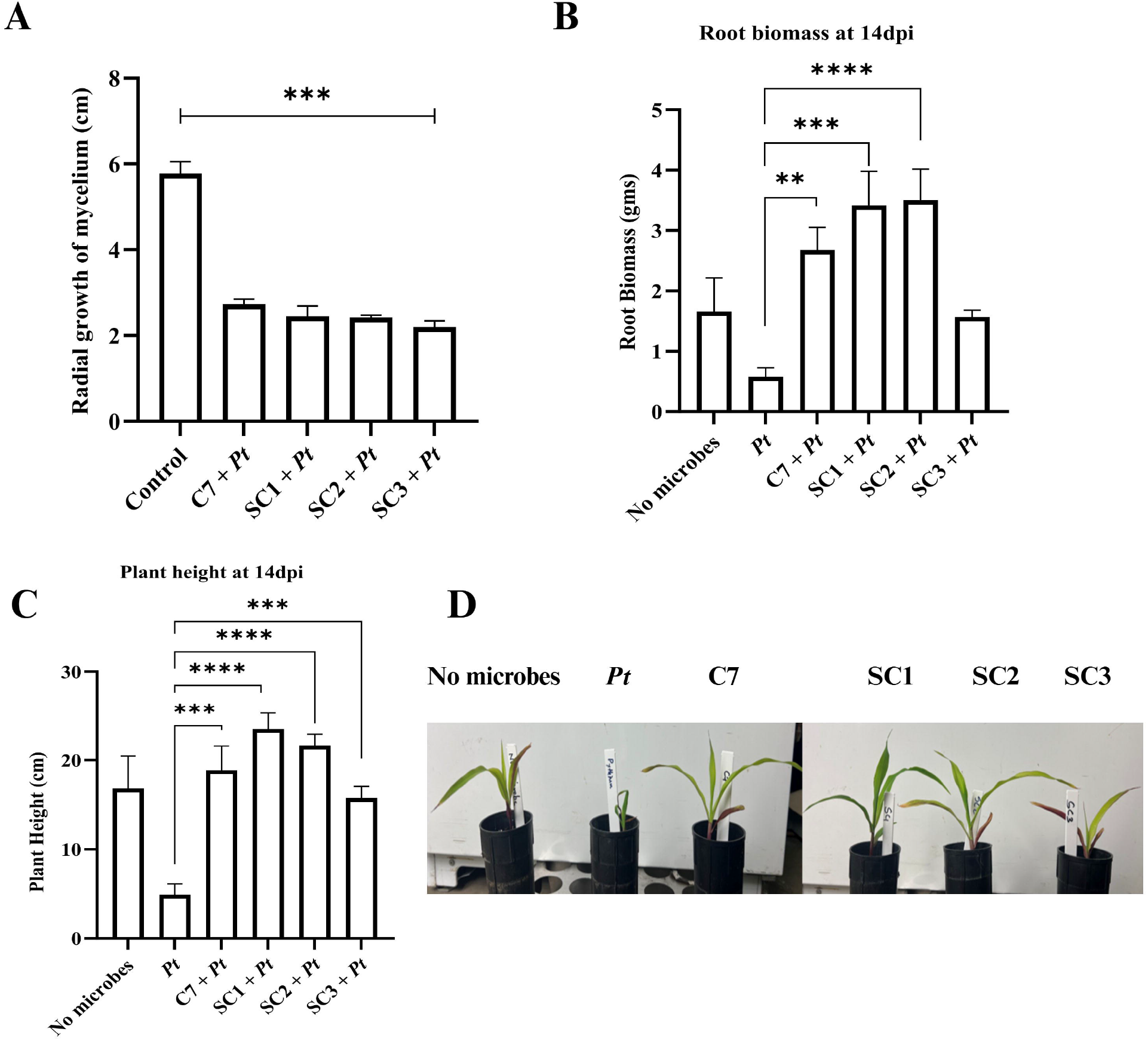
Antagonistic activity of sub-communities (SC1, SC2 and SC3) and C7 against corn oomycete. A) Measurement of radial growth of *Pythium torulosum* (*Pt*) in cm at 7 day post incubation. B) Pot assay of SC1, SC2 and SC3 and C7 community against the *Pythium torulosum* (*Pt*) where root biomass and C) and D) plant height was measured at 14 days after infecting (dpi) with biocontrol agents and pathogen. Three biological replications were maintained with each including technical replication (n = 5) for plate assay, and for pot assay (n = 7). Data are presented as mean ± Standard Deviation. For statistical analysis unpaired t-test with Welch’s correction.0’1+ 10.0.2) was used. ** <0.01, *** <0.001, **** <0.0001

### 3.7 Comparative analysis of auxin and siderophore production across the C7 community and subcommunities

We compared indole-3-acetic acid (IAA) synthesis and siderophore production among synthetic communities (SC1–SC3) and a previously tested community (C7). As shown in **Fig. 7A**, IAA production varied significantly among the communities and over time. SC2 exhibited the highest IAA levels, producing ∼45 µg/mL at day 2 and increasing to ∼65 µg/mL by day 7, indicating strong and sustained auxin synthesis. SC3 showed moderate IAA accumulation (∼20–25 µg/mL), while SC1 consistently produced the lowest levels (∼5–10 µg/mL). C7 maintained intermediate IAA levels (∼18 µg/mL) that remained stable between day 2 and day 7. In contrast, siderophore activity (**Fig. 7B**) was most pronounced in SC1 (0.55 cm, p = 0.01) and SC2 (0.50 cm, p = 0.015), followed by C7 (0.45 cm, p = 0.02). SC3 displayed the weakest response (0.35 cm, p = 0.093), which was not statistically significant. These findings indicate that community composition distinctly influences both siderophore and IAA production, where SC2 is the most efficient auxin producer, while SC1 and SC2 exhibit stronger siderophore activity, and C7 maintains intermediate levels of both traits.

**Figure 7.**
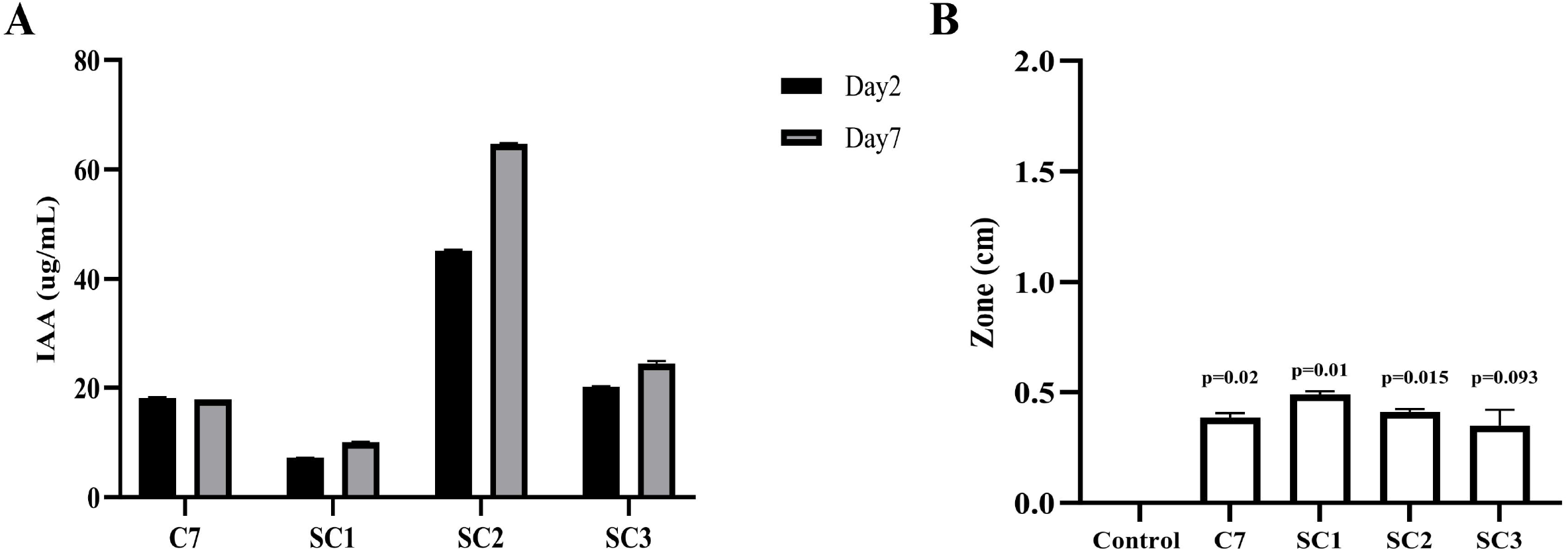
Assessment of IAA and siderophore production by communities (C7 and C8) and sub-communities (SC1, SC2 and SC3). (A) IAA production was quantified using the Salkowski reagent at 2 and 7 days of incubation. (B) Siderophore production was evaluated by measuring the diameter of the clear zone 7 days post-inoculation and compared with the control. Three biological replications were maintained with technical replications of (n =3). Data are presented as mean ± Standard Deviation. For statistical analysis, unpaired *t*-test with Welch’s correction.0’1+ 10.0.2) was used

## 4 Discussion

Although soil harbors immense microbial diversity that exceeds what is captured by 16S high-throughput sequencing, the functional outcomes of complex microbial species interactions and how plants recruit them to perform specific functions remain poorly understood. To investigate this, researchers have constructed synthetic bacterial communities, providing a simplified system to study microbial interactions and disease suppression. Early studies have used communities ranging from a few to several dozen strains (Lebeis et al., 2015; Yuan et al., 2018). For example, Niu et al. (2017) created a seven-member consortium to study maize rhizosphere colonization and pathogen suppression, while Armanhi et al. (2021) designed multi-strain communities to explore plant growth promotion in wheat. Such approaches have also been used to assemble sub-communities based on the performance of individual strains, allowing researchers to test whether specific combinations can achieve enhanced pathogen suppression compared with larger, more complex communities.

Our study builds on this framework by offering new insights into how individual strains, synthetic bacterial communities, and their subcommunities influence biocontrol activity against corn pathogens. We found that certain individual strains, communities, and subcommunities were significantly effective in preventing the growth of the oomycete pathogen, *P. torulosum*, but not so much for the fungal pathogens, as evident from the plate assays. Since the plate assays did not involve physical contact between the pathogen and the biocontrol agents, we suspect that secreted compounds may exhibit antagonistic activity against the pathogen. Furthermore, the compounds produced by these microbes may target the cell wall of the oomycete pathogen more effectively than the cell wall of the fungal pathogens. Oomycetes have cell walls that are primarily composed of cellulose, unlike fungal cell walls, which are mainly composed of chitin. This fundamental compositional difference could allow compounds targeting cellulose synthesis in oomycetes. For example, the cellulose synthase inhibitor dichlobenil disrupts oomycete growth and infection by *Phytophthora infestans*, while generally being ineffective against true fungi that lack cellulose in their cell walls (Grenville-Briggs et al. 2008), highlighting cellulose as a unique and vulnerable target specific to oomycetes. Apart from antibiosis, induced systemic resistance could also play a role in protecting the plant host from the pathogen (Gómez-Lama Cabanás et al. 2014). However, further investigations will be needed to understand the mode of action of these biocontrol agents.

Importantly, our results also highlight how extending or simplifying specific consortium members can alter the biocontrol phenotype. For instance, the extension of C7 with strain S8 did not lead to further improvements in plant growth or disease protection. This result was supported by the population profiling, which revealed that S8 failed to persist within the community. This outcome suggests that C7’s ecological stability and resilience prevented the integration of the added strain, a phenomenon consistent with priority effects and niche preemption in root-associated microbiomes. Similar patterns have been observed in synthetic communities of *Arabidopsis*, where Carlström et al. (2019) demonstrated that Alphaproteobacteria introduced at later stages were suppressed by pre-established communities, resulting in significant negative effects on Betaproteobacteria and Actinobacteria. Likewise, Niu et al. (2017) demonstrated that the removal of *Enterobacter cloacae* destabilized a maize root SynCom, allowing *Clavibacter pusillum* to take over, highlighting the role of keystone species in maintaining community integrity. Together, these findings reinforce that resident or native microbial consortia often resist colonization by external strains and that invasion success is strongly shaped by community history and assembly order (Gschwend et al., 2021).

Furthermore, comparisons between individual strains and the community also revealed trait-specific differences. While C7 provided better shoot height recovery under pathogen stress than any single strain, certain isolates, including S8, outperformed the community in promoting root biomass. These findings are consistent with previous work in corn, where Niu et al. (2017) reported that certain individual isolates showed stronger effects on specific plant traits, such as root growth promotion, while the synthetic community as a whole produced more consistent effects across different plant growth parameters. Taken together, this suggests that although single strains may excel in specific functional niches, multi-strain consortia buffer variability and contribute to more stable outcomes at the whole-plant level.

Several biocontrol agents also promote plant growth through diverse biochemical and molecular mechanisms. For instance, *Bacillus subtilis* MBI600 was recently shown to enhance tomato shoot and root development, in part by activating auxin-related genes (SiPin6 and SiLax4), while also suppressing major soilborne pathogens such as *Rhizoctonia solani, Pythium ultimum*, and *Fusarium oxysporum* f. sp. *Radicis-lycopersici* (Samaras et al., 2021). Similarly, siderophore-producing strains such as *Pseudomonas aeruginosa* F2 and *P. fluorescens* JY3 not only restricted the growth of *F. oxysporum* and *R. solani* through iron competition but also stimulated wheat growth. Bioformulations reduced damping-off by up to 87.5% and enhanced shoot and root biomass under greenhouse conditions. These examples demonstrate how biocontrol strains offer dual benefits by simultaneously promoting plant growth and suppressing pathogens (Abo-Zaid et al., 2023). In our study, the free-living nitrogen-fixing bacterium *Azotobacter vinelandii* (S8) demonstrated strong antagonism against *Pythium torulosum* and moderate suppression of *Fusarium* spp., while also enhancing plant height and root biomass under pathogen pressure. These results confirm the dual role of S8 as both a plant growth–promoting bacterium and a biocontrol agent, consistent with previous reports on the contribution of *Azotobacter* species to biological nitrogen fixation and disease suppression (Aasfar et al., 2021). In this context, our evaluation of plant growth–promoting traits, such as IAA and siderophore production, further illustrates the importance of functional specialization and community-level buffering. Individual strains exhibited wide variation, with S4 being a strong auxin producer and S5 excelling in siderophore production, while others, such as S7 and S8, maintained consistently low metabolite output. In contrast, both C7 and C8 communities produced intermediate and stable levels of IAA and siderophores, underscoring the stabilizing effect of microbial interactions on key community-level functions. Such balancing has also been observed in synthetic microbiome studies, where community performance often exceeds the variability of individual members, providing functional reliability even if populations of single strains fluctuate (Herrera Paredes et al., 2018).

Most notably, rationally designed subcommunities (SC1 and SC2) surpassed the performance of C7 in promoting plant growth and root biomass under pathogen pressure. SC1 and SC2 likely succeeded because they contained fewer, but complementary strains, which minimized negative cross-talk while maximizing cooperative interactions. For instance, SC2 combined high IAA-producing members, which explains its superior auxin synthesis over time, while SC1 and SC2 retained higher siderophore activity, reflecting a more substantial iron acquisition potential. By contrast, SC3 did not perform as strongly across these traits, suggesting that its strain composition may have lacked sufficient complementarity or increasingly incompatible interactions among the microbes. These results suggest that trait complementarity, rather than taxonomic diversity or community size, is key to the design of effective biocontrol consortia. Similar conclusions have been reported in studies showing that smaller, functionally compatible synthetic communities can outperform more complex mixtures by avoiding antagonistic interactions and ensuring consistent functional output (Vacheron et al., 2013; Camarillo et al., 2023).

Our findings highlight the importance of developing next-generation biocontrol technologies in agriculture. The superior performance of streamlined subcommunities (SC1 and SC2) demonstrates that functionally complementary but minimal consortia can outperform larger mixtures. Such simplified inoculants are easier to formulate and more likely to remain stable under field conditions, making them attractive for large-scale deployment. The inability of S8 to integrate into C8 underscores the importance of community assembly history, priority effects, and niche occupation, which must be considered when designing inoculants for diverse soils and cropping systems. Equally important, trait-guided community design such as combining high auxin producers with strong siderophore producers offers a rational strategy for creating inoculants that both protect against pathogens and promote plant growth. This dual functionality mirrors successful examples from other systems, including *Bacillus subtilis* MBI600, which enhances tomato growth while suppressing soilborne pathogens (Samaras et al., 2021), and siderophore-producing *Pseudomonas* strains that reduce damping-off in wheat while boosting biomass (Abo-Zaid et al., 2023). Building on these insights, microbial consortia tailored for corn could be delivered as seed coatings, soil drenches, or granular formulations, providing sustainable alternatives to chemical pesticides and fertilizers.

Ultimately, rationally designed microbial consortia have the potential to improve yield stability, reduce chemical inputs, and enhance resilience to soilborne diseases. Future work should evaluate these subcommunities across diverse soils and environments, refine formulations for field durability, and leverage synthetic biology to engineer more predictable and synergistic interactions (Alori and Babalola, 2018).

## 5 Conclusion

This study demonstrates that microbial community performance is driven more by functional complementarity than by taxonomic diversity or community size. Rationally designed subcommunities, particularly SC1 and SC2, consistently suppressed *Pythium* and promoted corn growth more effectively than larger, less coordinated mixtures. These findings highlight the potential of minimal, functionally complementary consortia as scalable and reliable biocontrol solutions. Future efforts should focus on testing these streamlined inoculants under field conditions and refining trait-guided approaches for designing robust microbial products that support sustainable crop production.

## Author Contributions

**PK -** Writing – review and editing, Methodology, Writing – original draft, Conceptualization, Investigation, Formal analysis, Validation. **DS** - Formal analysis, Writing - review and editing. **DK** - Conceptualization, Writing – review and editing.

## Acknowledgements

We thank Dean Malvick and Crystal Floyd for sharing corn fungal and oomycete strains with us. We also thank Mia Copeland and Ashish Ranjan for sharing protocols for testing siderophore and IAA activities. The use of any trade, firm, or corporation names in this publication is for the information and convenience of the reader. Such use does not constitute an official endorsement or approval by the US Department of Agriculture or the Agricultural Research Service of any product or service to the exclusion of others that may be suitable. The findings and conclusions in this publication are those of the authors and should not be construed to represent any official USDA or US Government determination or policy. USDA is an equal opportunity provider and employer.

## Generative AI statement

The authors declare that no Gen AI was used in the creation of this manuscript.

